# Abnormal enteric nervous system and motor activity in the ganglionic proximal bowel of Hirschsprung’s disease

**DOI:** 10.1101/2023.03.08.531750

**Authors:** Brian S. Edwards, Emma S. Stiglitz, Brian M. Davis, Kristen M. Smith-Edwards

**Author notes:** Corresponding author: Kristen M. Smith-Edwards Department of Physiology and Biomedical Engineering, Mayo Clinic, 200 First Street SW, Guggenheim Building 15-11C, Rochester, MN 55905.

## Abstract

Hirschsprung’s disease (HSCR) is a congenital defect in which the enteric nervous system (ENS) does not develop in the distal bowel, requiring surgical removal of the portions of bowel without ENS ganglia (‘aganglionic’) and reattachment of the ‘normal’ proximal bowel with ENS ganglia. Unfortunately, many HSCR patients have persistent dysmotility (e.g., constipation, incontinence) and enterocolitis after surgery, suggesting that the remaining bowel is not normal despite having ENS ganglia. Anatomical and neurochemical alterations have been observed in the ENS-innervated proximal bowel from HSCR patients and mice, but no studies have recorded ENS activity to define the circuit mechanisms underlying post-surgical HSCR dysfunction. Here, we generated a HSCR mouse model with a genetically-encoded calcium indicator to map the ENS connectome in the proximal colon. We identified abnormal spontaneous and synaptic ENS activity in proximal colons from GCaMP-Ednrb^-/-^ mice with HSCR that corresponded to motor dysfunction. Many HSCR-associated defects were also observed in GCaMP-Ednrb^+/-^ mice, despite complete ENS innervation. Results suggest that functional abnormalities in the ENS-innervated bowel contribute to post-surgical bowel complications in HSCR patients, and HSCR-related mutations that do not cause aganglionosis may cause chronic colon dysfunction in patients without a HSCR diagnosis.

## INTRODUCTION

Hirschsprung’s disease (HSCR) is a congenital defect caused by a number of known gene mutations (e.g., *RET, EDNRB*) and results in distal bowel without ganglia of the enteric nervous system (ENS) (1) required for the coordinated contractions and relaxations of smooth muscle that move contents along the gastrointestinal tract (2, 3). In HSCR, the absence of neurally-mediated motor patterns in the aganglionic distal regions, where ENS ganglia are missing, leads to tonic constriction and the inability to move fecal contents, thereby causing functional obstruction. The portion of bowel proximal to the constricted, aganglionic region becomes extremely dilated and is known as ‘megacolon’ (1). This potentially life-threatening condition necessitates the immediate surgical removal of aganglionic bowel and reattachment of the ‘normal’ ganglionic bowel that is innervated by the ENS. Unfortunately, even after surgical treatment many children with HSCR continue to suffer from bowel dysmotility (e.g., constipation, incontinence) and enterocolitis (1, 4–10), suggesting that the remaining bowel is not normal despite the presence of ENS ganglia. To better treat post-surgical bowel complications in HSCR patients, we need to know the extent that HSCR-mutations alter the structure and function of the ENS in the remaining portions of bowel that have been presumed to be normal.

The ENS consists of a heterogeneous population of up to 20 different neuron types distinguished by morphology, location, electophysiological properties, neurotransmitters, and connectivity (11), which communicate with one another in a tightly coordinated manner to properly execute bowel functions. Myenteric neurons of the ENS are located between the circular and longitudinal muscle layers and form complex circuits that generate reflexive motor patterns necessary for propulsive motility. Intrinsic primary afferent neurons located within myenteric ganglia detect stretch and other mechanical forces to activate interneurons and motor neurons that are organized into polarized pathways (12, 13). Ascending pathways within the ENS travel orally and contain interneurons and excitatory motor neurons that release acetylcholine (and other excitatory neurotransmitters) to contract smooth muscle, whereas descending pathways travel anally and contain interneurons and inhibitory motor neurons that release nitric oxide (and other inhibitory neurotransmitters) to relax smooth muscle (11, 14, 15). In addition to myenteric neurons, motor patterns are regulated by interstitial cells of Cajal (ICC), also known as the ‘pacemakers’ of the gut (16). ICC form electrically-coupled networks that are intimitately associated with smooth muscle cells and the ENS (17–20). Therefore dysregulation of ICC activity, due to intrinsic or neurally-mediated changes, can also have major implications on motility behavior.

A growing body of evidence from HSCR patients and mouse models has reported anatomical and neurochemical alterations in the ENS-innervated proximal bowel. In addition to smaller-sized ganglia with reduced numbers of neurons, evidence from patients and mouse models indicates that the ganglionic bowel has increased nitrergic myenteric neurons and decreased cholinergic myenteric neurons (21–25), suggesting an imbalance in the ascending excitatory and descending inhibitory circuits required for propulsion. Changes in the density and phenotype of ICC have also been shown in ENS-innervated samples from HSCR patients and mouse models (26–28) that may also influence motility. But despite the numerous examples of anatomical abnormalities in HSCR patients and mice and reports of altered motor patterns in HSCR mouse models (23, 24), no studies have directly recorded changes in cellular activity to define the ENS circuit mechanisms underlying HSCR-associated dysmotility. This information is critical for the development of targeted strategies to improve post-surgical bowel complications in HSCR.

Here, we generated the first HSCR mouse model that also expresses a genetically-encoded calcium indicator to map and identify functional defects in the ENS connectome of the HSCR proximal colon. We bred an E2a-GCaMP mouse line that we have previously used to study the ENS and ICC in healthy, adult mice (29–31), to the well-characterized and physiologically-relevant HSCR mouse model with loss-of-function mutations in the gene for endothelin receptor B (*EDNRB*). We chose this particular HSCR model because *EDNRB* mutations contribute to ∼5% of familial HSCR cases (32–37), and unlike mice with other HSCR-associated mutations, *EDNRB* mutant mice phenocopy distal bowel aganglionosis observed in HSCR patients and survive up to 4 weeks (38). Spontaneous and synaptically-evoked activity in myenteric neurons and ICC, as well as colonic motor patterns, were recorded in *ex vivo* colon preparations from GCaMP-Ednrb^+/+^ mice (i.e., wildtype mice expressing GCaMP), GCaMP-Ednrb^+/-^ mice (i.e., mice lacking one copy of the *EDNRB* gene and expressing GCaMP), and GCaMP-Ednrb^-/-^ mice (i.e., mice lacking both copies of the *EDNRB* gene and expressing GCaMP) at post-natal day 15 (P15). We identified major differences in the organization, neurochemical phenotype, and activity of myenteric neurons that corresponded to abnormal motility behavior in proximal colons from P15 GCaMP-Ednrb^-/-^ mice that have HSCR-like phenotype. Surprisingly, many of these HSCR-associated defects were also observed in proximal colons from P15 and adult GCaMP-Ednrb^+/-^mice, despite complete innervation by ENS ganglia (i.e., no overt HSCR-like aganglionosis). In conclusion, our results revealed several defects in the ENS of the proximal colon in mice with *EDNRB* mutations that likely contribute to post-surgical bowel issues in HSCR patients. Moreover, HSCR-related mutations that do not cause aganglionosis may produce chronic colon dysfunction in patients without a HSCR diagnosis.

## RESULTS

### Generation and characterization of GCaMP-Ednrb model for Hirschsprung’s Disease

The goal of this study was to test the hypothesis that long-term bowel complications in HSCR after surgical treatment is due to ENS dysfunction in the proximal bowel that has been presumed to be normal based on the presence of ENS ganglia. Towards this goal, we bred mice heterozygous for the loss-of-function mutation *EDNRB* (Ednrb^tm1Ywa^) to our established E2a-GCaMP mouse line (29–31), generating an HSCR mouse model (referred to as GCaMP-Ednrb) with a genetically-encoded calcium indicator to record activity from the ENS and ICC (**Figure 1A**). Interestingly, most GCaMP-Ednrb^-/-^ offspring were males (80% compared to 64% and 58% in GCaMP-Ednrb^+/-^ and GCaMP-Ednrb^+/+^ mice, respectively), thus exhibiting remarkable similarity to the clinical prevelance of HSCR in males (4:1 male to female ratio) (39, 40). Consistent with previous reports (38), GCaMP-Ednrb^-/-^ mice with HSCR died prematurely, typically in the third week of life (range: 8-24 days). Because the ENS undergoes rapid, post-natal developmental changes during this time (41–43), we performed experiments in each genotype at post-natal day 15 (P15), a timepoint when the major motor patterns have developed (24) but before the majority of mice with HSCR became septic and died.

**Figure 1.**
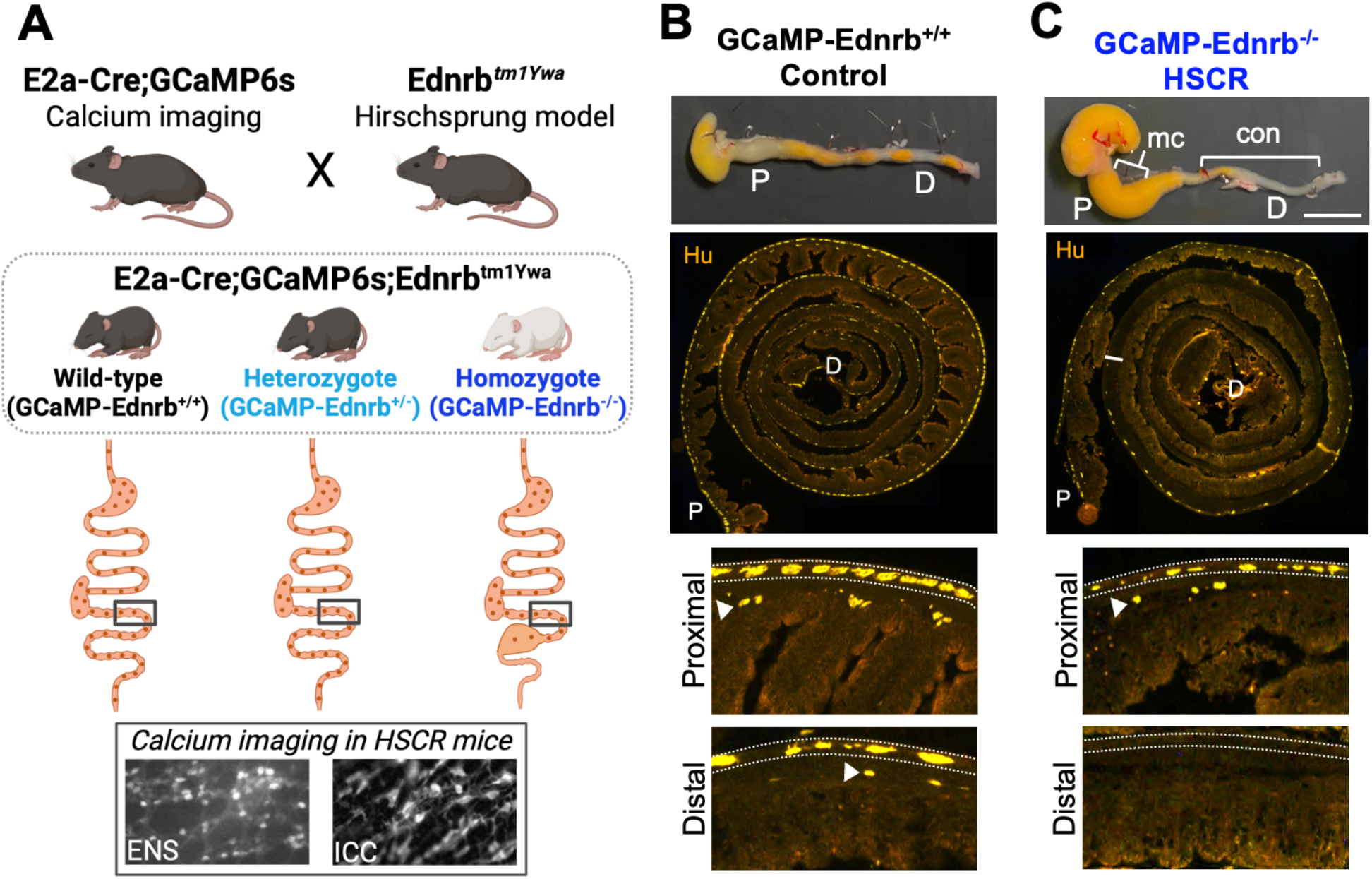
Characterization of the first GCaMP-HSCR transgenic mouse model. (A) Schematic overview of the generation and use of a new GCaMP model to study HSCR. Top, we bred our well-established E2a-Cre;GCaMP mouse line to mice with heterozygous knock-out mutations in the gene for endothelin receptor B (Ednrb) to generate a novel HSCR model that also has the genetically encoded calcium indicator, GCaMP6s, expressed in neuronal and non-neuronal cell types. The gray dotted box indicates the genotypes of experimental mice used in the present studies. Below each mouse and genotype are cartoon depictions of the gastrointestinal (GI) tract and innervation by ENS ganglia (tan dots) that show normal macroscopic appearance and ENS innervation of the GI tract in GCaMP-Ednrb^+/+^ and GCaMP-Ednrb^+/-^ mice, whereas GCaMP-Ednrb^-/-^ mice have an aganglionic, constricted distal bowel with megacolon in the adjacent, proximal segment; note that the rest of the GI tract appears normal. Thus, the generation of the GCaMP-Ednrb mouse model allowed us, for the first time, to record activity in the ENS and interstitial cells of cajal (ICC) to directly test whether the ENS-innervated bowel, that remains after surgery, functions normally in HSCR; example still images of GCaMP signal during recordings of each each cell type are shown at the bottom. (B-C) Top, example colons from GCaMP-Ednrb^+/+^ (B) and GCaMP-Ednrb^-/-^ (C) mice, where the constricted bowel, dilated megacolon, and back-up of fecal contents are visible. Middle, colon “swiss” roll preparations from GCaMP-Ednrb^+/+^ (B) and GCaMP-Ednrb^-/-^ (C) mice were stained with the pan-neuronal marker, Hu. Note the absence of Hu+ ganglia in the distal colon in C (white line indicates transition from aganglionic to ganglionic colon). Bottom, close-up images from proximal and distal regions of colons from GCaMP-Ednrb^+/+^ (B) and GCaMP-Ednrb^-/-^ (C) mice that clearly show distal colon aganglionosis in GCaMP-Ednrb^-/-^ mice; dotted lines indicate the myenteric plexus where myenteric neurons reside, and arrows indicate the submucosal plexus where submucosal neurons reside. P, proximal colon; D, distal colon; mc, megacolon; con, constricted.

As expected, colons from GCaMP-Ednrb^+/+^ and GCaMP-Ednrb^+/-^ mice were completely innervated by the ENS and appeared normal macroscopically (**Figure 1B**), whereas all GCaMP-Ednrb^-/-^ mice exhibited aganglionosis, as well as the hallmark abnormalities in colon shape observed in HSCR patients and mouse models (1, 38). A constricted distal bowel that correlated to the aganglionic segment and a dilated ‘megacolon’ was observed proximal to the constricted region in GCaMP-Ednrb^-/-^ mice with HSCR (**Figure 1C**). The extent of aganglionosis in GCaMP-Ednrb^-/-^ mice varied; 47% (7/15) of mice exhibited aganglionosis that was restricted to the distal half of the colon, 33% (5/15) of mice had aganglionosis that extending into more proximal regions of the colon, and 20% (3/15) of mice had total colonic aganglionosis (Supplemental Table 1). The three mice with total colonic aganglionosis were excluded from the experiments for this study since it was not possible to record ENS activity in the same region of proximal colon recorded in all other mice.

### HSCR-associated differences in ENS organization and neurochemical properties in proximal colon

A closer inspection of the ENS in whole-mount preparations revealed significant abnormalities in the organization and neurochemical identity of myenteric neurons in proximal colons from GCaMP-Ednrb HSCR mice. We stained tissue with the pan-neuronal marker HuC/D, as well as nitric oxide synthase (NOS) or calretinin (CalR) to broadly label inhibitory or excitatory enteric neurons, respectively (11) (**Figure 2A-B**). The proximal colons from GCaMP-Ednrb^-/-^ mice had significantly smaller ganglia (**Figure 2C**) and fewer myenteric neurons per field of view (**Figure 2D**) compared to the same region in GCaMP-Ednrb^+/+^ and -Ednrb^+/-^ mice, indicating that the ENS-innervated proximal colon in HSCR mice is hypoganglionated. Surprisingly, the proximal colons from GCaMP-Ednrb^+/-^ mice also had significantly smaller ganglia and lower number of myenteric neurons compared to Ednrb^+/+^ wild-type littermates, suggesting that although heterozygote mutations in *EDNRB* did not produce HSCR-like aganglionosis in mice, these potentially ‘subclinical’ mutations led to detectable changes in the development and organization of the ENS. The large majority of myenteric neurons in proximal colons from GCaMP-Ednrb^-/-^ mice expressed NOS, with an average percentage of NOS+ myenteric neurons that was significantly greater compared to both GCaMP-Ednrb^+/+^ and GCaMP-Ednrb^+/-^ mice (**Figure 2E**). It was also noted that GCaMP-Ednrb^+/-^ mice had a significantly greater percentage of NOS+ neurons compared to GCaMP-Ednrb^+/+^ mice (**Figure 2E**). The increase in NOS+ myenteric neurons in GCaMP-Ednrb^-/-^ proximal colons was accompanied by a decrease in CalR+ myenteric neurons compared to GCaMP-Ednrb^+/-^ and almost reached significance when compared to GCaMP-Ednrb^+/+^ mice (**Figure 2F**).

**Figure 2.**
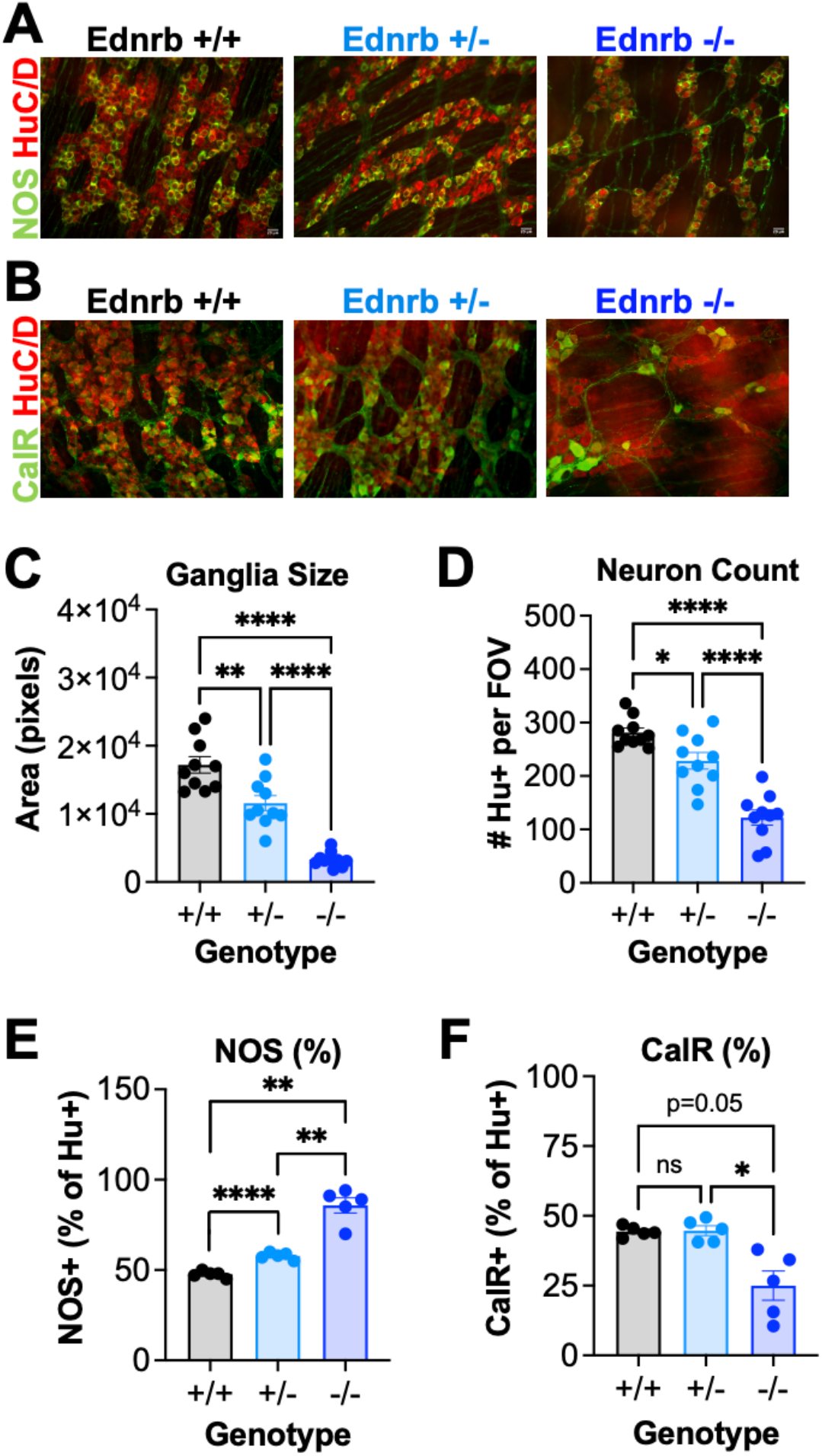
Altered ENS organization and neurochemical expression in mice with Ednrb mutations. (A) Example images of proximal colon tissue from each genotype stained with nitric oxide synthase (NOS; green) and Hu (red). (B) Example images of proximal colon tissue from each genotype stained with calretinin (CalR; green) and Hu (red). (C) On average, proximal colons from mice with homozygous (hom, -/-) Ednrb mutations had significantly smaller ganglia by area compared to mice with heterozygous (het, +/-) Ednrb mutations and wild-type littermate mice (p<0.0001 for both), and mice with het (+/-) Ednrb mutations had significantly smaller ganglia compared to wild-type littermate mice (p=0.0094; n=10 mice, Welch’s ANOVA with Dunnet’s test for multiple comparisons). (D) The average number of Hu+ myenteric neurons per field of view was also significantly lower in colons from mice with hom (-/-) Ednrb mutations compared to mice with het (+/-) Ednrb mutations and wild-type littermate mice (p<0.0001); and again, significantly fewer Hu+ myenteric neurons were found in colons from mice with het (+/-) Ednrb mutations compared to wild-type littermate controls (p=0.0213, n=10; one-way ANOVA with Tukey’s test for multiple comparisons). (E) The average percentages of NOS+ myenteric neurons in mice with het (+/-) and hom (-/-) Ednrb mutations were significantly higher compared to wild-type littermate mice (p<0.001 and p=0.0023, respectively, n=10; Welch’s ANOVA with Dunnet’s test for multiple comparisons), and NOS+ percentages were significantly higher in mice with hom (-/-) Ednrb mutations compared to mice with het (+/-) Ednrb mutations (p=0.0073). (F) The average percentage of CalR+ myenteric neurons in mice with hom (-/-) Ednrb mutations was decreased compared to the other genotypes, but this was only statistically significant compared to mice with het (+/-) mutations, which were no different compared to wild-type (+/+) mice (p=0.0430, n=10, Welch’s ANOVA with Dunnet’s test for multiple comparisons).

### Abnormalities in spontaneous ENS, ICC, and motor activity in ‘HSCR’ proximal colon

In healthy adult GCaMP mice, myenteric neurons exhibit ongoing activity that contribute to spontaneous motility in the proximal colon (29), and we found that this was also true in P15 GCaMP-Ednrb mice (**Figure 3A**). However, compared to GCaMP-Ednrb^+/+^ control mice, the percentage of spontaneously active myenteric neurons in proximal colons from GCaMP-Ednrb^-/-^ and GCaMP-Ednrb^+/-^ mice was significantly decreased (**Figure 3B**), indicating functional abnormalities in the proximal colon even in mice lacking a single HSCR-associated allele and no evidence of an aganglionic distal colon (i.e., in GCaMP-Ednrb^+/-^ mice).

**Figure 3.**
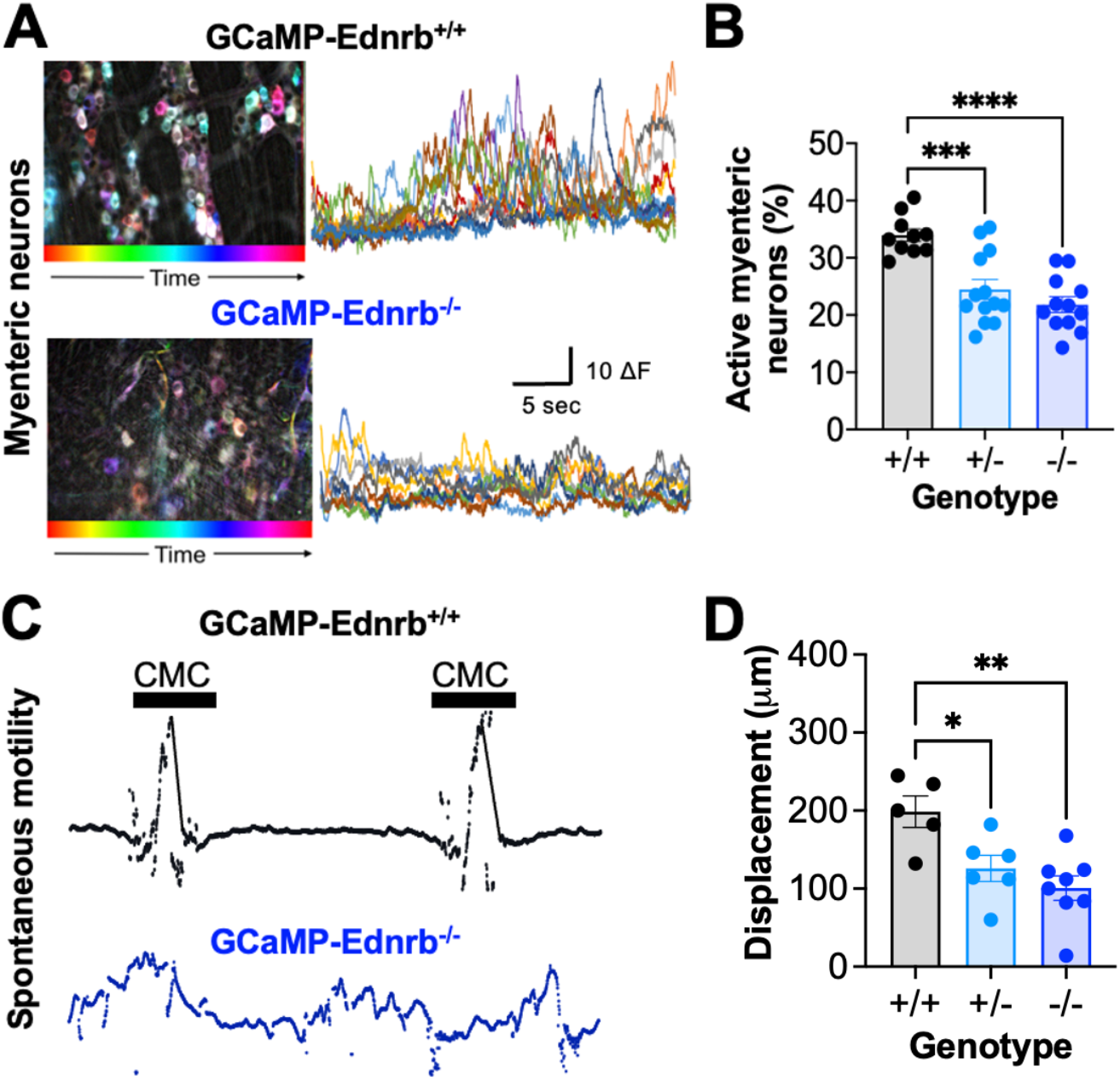
Differences in baseline myenteric neuron activity and spontaneous motor patterns in mice with Ednrb mutations. (A) Time-lapse color coded images (left) and traces (right) of spontaneous GCaMP activity from individual myenteric neurons in proximal regions of colons from GCaMP-Ednrb^+/+^ wild-type mice (top) and GCaMP-Ednrb^-/-^ mice that have HSCR (bottom). Images were made in ImageJ using the “Time-lapse color coder” plugin, where each pixel is assigned a color based on when it reached maximum fluorescence intensity. (B) Graph of the average percentage of myenteric neurons in a field of view with spontaneous activity in a 30-sec period; significant decreases were observed in GCaMP-Ednrb^-/-^ and GCaMP-Ednrb^+/-^ mice compared to GCaMP-Ednrb^+/+^ (p<0.0001 and p=0.0006, respectively, n=12, one-way ANOVA with Tukey’s test for multiple comparisons). (C) Example traces of displacement within the imaging field showing motility patterns in the proximal colon from GCaMP-Ednrb^+/+^ (wild-type) (top, black) and GCaMP-Ednrb^-/-^ (bottom, blue) mice; note the distinct appearance of colon motor complexes (CMC) indicated by black bars in the black trace, whereas the motor patterns exhibit in the blue trace are uncoordinated and do not resemble normal CMCs. (D) Graph of the maximum displacement in proximal colon imaging fields reveal significant decreases in GCaMP-Ednrb^-/-^ and GCaMP-Ednrb^+/-^ mice compared to GCaMP-Ednrb^+/+^ mice (p=0.0032 and p=0.0359, respectively, n=5-8 mice, one-way ANOVA with Tukey’s test for multiple comparisons).

To see if changes in myenteric neuron activity correlated to abnormal motor activity, we compared spontaneous motility behavior in isolated colons from GCaMP-Ednrb mice by collecting 5 consecutive 3-min calcium imaging recordings and plotting tissue displacement shown to be correlated to changes in tension (31). In full length, full thickness colon preparations from P15 GCaMP-Ednrb^+/+^ and GCaMP-Ednrb^+/-^ mice, colon motor complexes (CMC) were regularly generated in the proximal colon every 2-4 min, and each CMC was separated by a period of time with relatively little motor activity (**Figure 3C**, black traces). However, the CMCs produced in GCaMP-Ednrb^+/-^ mice were significantly lower in magnitude compared to GCaMP-Ednrb^+/+^ mice (**Figure 3D**). By contrast, definitive CMCs were not observed in preparations from GCaMP-Ednrb^-/-^ mice; instead there were continuous contractions that periodically appeared to grow in amplitude every 1-2 min (**Figure 3C**, blue traces). Although the frequency and duration of these contractions were increased compared to the CMCs in GCaMP-Ednrb^+/+^ mice, the magnitude was significantly decreased (**Figure 3D**), suggesting that the rhythmic contractions produced in proximal colons from mice with HSCR are not productive or efficient for propulsion.

Interstitial cells of Cajal (ICC) are another major cell type involved in producing rhythmic motor patterns. Specifically, ICCs in the submucosal plexus (ICC-SM) of the colon produce slow waves of depolarization responsible for “ripple” contractions (20), and we recorded and compared ICC activity in proximal and distal colon (**Figure 4A**). There were no significant differences in the frequency of ICC-SM oscillations in proximal colon across genotypes, but ICC-SM oscillations in the distal colon were significantly lower in frequency in both GCaMP-Ednrb^-/-^ and GCaMP-Ednrb^+/-^ mice compared to controls (**Figure 4B**). As shown in **Figure 4C**, the frequency of ICC-SM oscillations was higher in distal colon compared to proximal colon from GCaMP-Ednrb^+/+^ and GCaMP-Ednrb^+/-^ mice (also true in the healthy adult colon, unpublished data, Smith-Edwards). These differences demonstrate that a frequency gradient is normally established across the length of colon that influences the directionality and velocity of slow waves produced by ICC networks (17) and therefore also affects ripple contractions. However, in GCaMP-Ednrb^-/-^ mice, the ICC-SM oscillation frequencies were not significantly different between proximal and distal colon (**Figure 4C**), indicating a lack of or reduced frequency gradient in most GCaMP-Ednrb^-/-^ mice. We then measured the frequency gradient in each colon preparation by calculating the difference in oscillation frequency at each region (e.g., frequency in distal colon minus frequency in proximal colon). The frequency gradient in preparations from GCaMP-Ednrb^+/-^ and GCaMP-Ednrb^-/-^ mice were significantly lower than GCaMP-Ednrb^+/+^ control mice (**Figure 4D**). The subtle changes in ICC slow wave properties may partially contribute to the dysrhythmic spontaneous motility observed in Ednrb mutant mice by disrupting the ability for ICC to organize slow wave activity across large networks.

**Figure 4.**
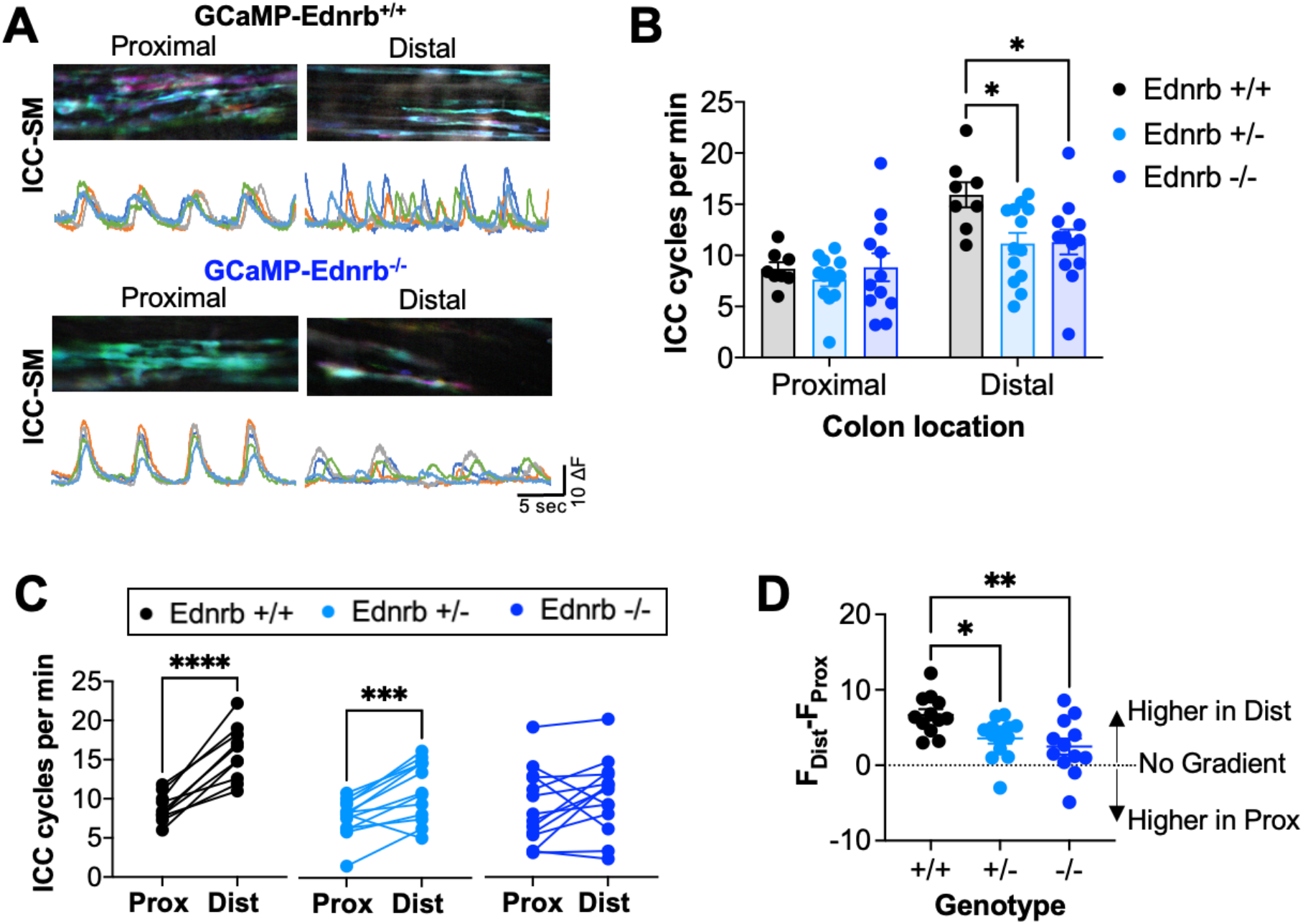
Minimal changes in ICC slow wave activity. (A-B) Example ICC slow wave calcium recordings in proximal and distal colon regions from GCaMP-Ednrb^+/+^ mice (A) and GCaMP-Ednrb^-/-^ mice (B); time-lapse color coded images during a single oscillation shown on top and GCaMP traces of individual ICC shown on bottom. (C) Normal ICC slow waves result from a gradient, with significantly lower oscillation frequencies in proximal compared to distal regions (black data points; p<0.0001 in GCaMP-Ednrb^+/+^, n=12, paired Student’s t-test), and ICC oscillation frequencies in GCaMP-Ednrb^+/-^ mice were also significantly lower in proximal colon (light blue data points; p=0.0002, n=14, paired Student’s t-test). However, there were no significant differences between proximal and distal ICC oscillation frequencies in GCaMP-Ednrb^-/-^ mice. (D) Plot of ICC frequency gradients (measured as frequency in proximal colon minus frequency in distal colon) shows significantly lower gradients (i.e., closer to zero) were recorded from GCaMP-Ednrb^-/-^ and GCaMP^+/-^ mice compared to GCaMP-Ednrb^+/+^ mice (p=0.0039 and p=0.0290, respectively, n=12-14, one-way ANOVA with Tukey’s test for multiple comparisons).

### HSCR-associated defects in ENS circuit connectivity and evoked contractions

Normal colon motility and propulsion of fecal contents requires coordinated, multi-synaptic activity in ascending (oral-projecting) and descending (anal-projecting) myenteric neuron circuits. To assess synaptic connectivity within myenteric neuron circuits of the proximal colon in HSCR mice, we measured GCaMP responses to electrical stimulation 3 mm oral and anal to the field of view (**Figure 5A**), a distance at which most responses are blocked with hexamethonium and therefore represent synaptic activity rather than direct activation of fibers of passage (31). Myenteric neurons were characterized based on whether they responded to oral stimulation, anal stimulation or both sites of stimulation, indicative of receiving descending, ascending, or bi-directional input, respectively (**Figure 5A**). Examples of responses from individual myenteric neurons in a field of view and their corresponding response types are shown in **Figure 5B**. GCaMP-Ednrb^-/-^ mice had a significantly greater percentage of myenteric neurons that specifically responded to oral stimulation compared to GCaMP-Ednrb^+/+^ mice (**Figure 5C**), indicating stronger descending input to myenteric neurons in the HSCR proximal colon. Additionally, both GCaMP-Ednrb^+/-^ and GCaMP-Ednrb^-/-^ mice had significantly lower percentages of myenteric neurons that specifically responded to anal stimulation compared to GCaMP-Ednrb^+/+^ mice, suggesting weaker ascending input to myenteric neurons in the proximal colon. Although less input from anal regions might have been expected in GCaMP-Ednrb^-/-^ mice because of their distal aganglionisis, the decrease observed in GCaMP-Ednrb^+/-^ mice suggest a dose-dependent effect of *EDNRB* gene deletion. That is, loss of even one copy of the EDNRB gene in mice can cause functional changes in synaptic connectivity without producing the more dramatic features of HSCR, such as aganglionosis.

**Figure 5.**
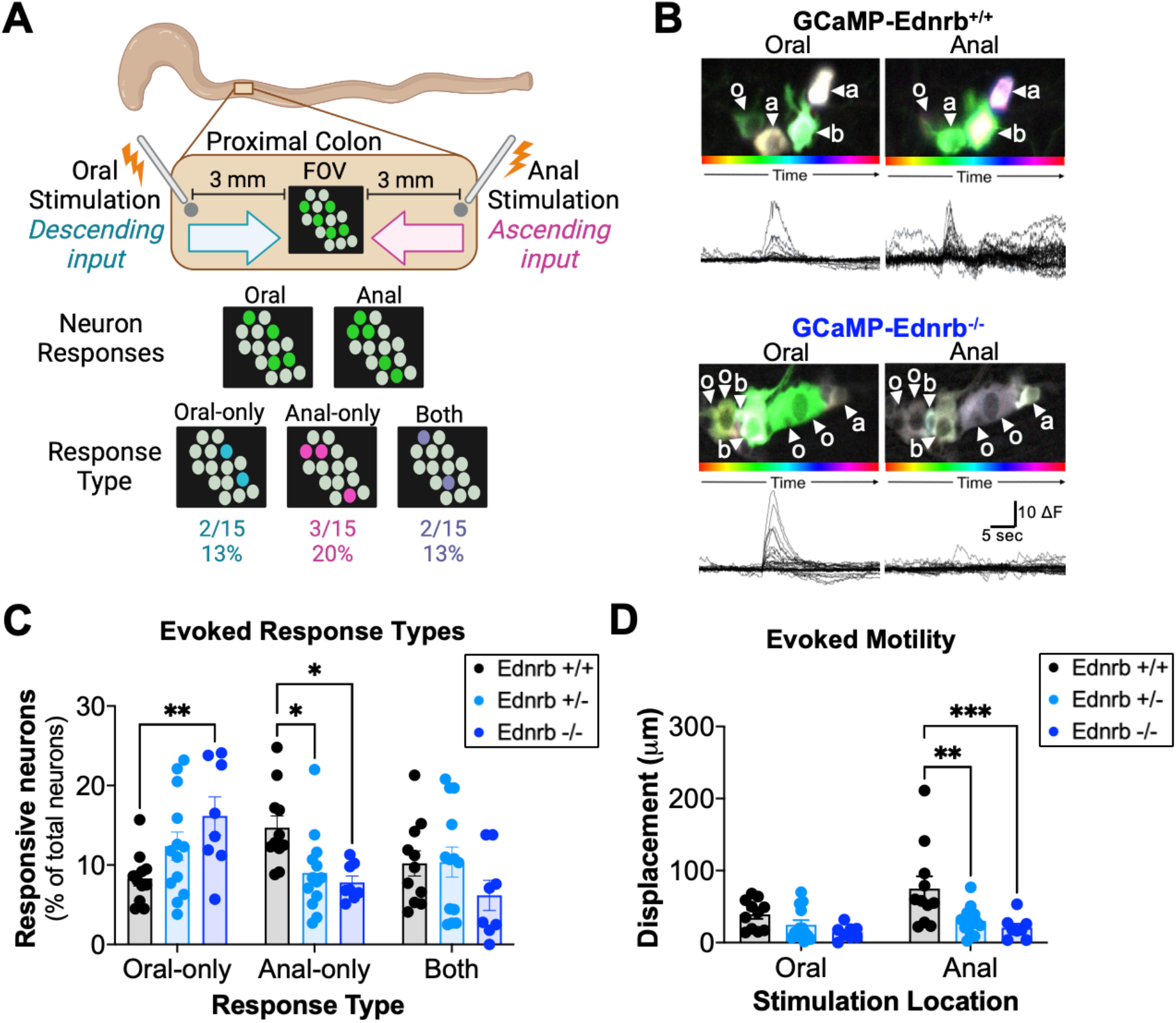
Alterations in myenteric neuron circuit connectivity and evoked contractions. (A) Experimental paradigm to measure the strength of ascending and descending neuronal circuits in the proximal colon, as previously published (cite). For each field of view (FOV), myenteric neuron and motility responses to oral and anal electrical stimulation were recorded. Results are reported as the percentage of neurons that responded only to oral stimulation, only to anal stimulation, or to both stimulation sites. (B) Example time-lapse color coded images and traces of GCaMP activity in response to electrical stimulation 3mm oral and anal to the imaging field in the proximal colon from GCaMP-Ednrb^+/+^ (top) and GCaMP-Ednrb^-/-^ mice (bottom). (C) Graph of the average percentages of each response type (oral-only, anal-only, or both) indicates a significant increase in myenteric neurons that only responded to oral stimulation (i.e., receive descending input) in GCaMP-Ednrb^-/-^ mice compared to GCaMP-Ednrb^+/+^ mice (p=0.0069), and significant decreases in myenteric neurons that only respond to anal stimulation (i.e., receive ascending input) in both GCaMP-Ednrb^-/-^ (p=0.0219) and -Ednrb^+/-^ mice (p=0.0349) compared to GCaMP-Ednrb^+/+^ mice; there were no differences in the percentage of myenteric neurons that responded to both stimulation sites (i.e., receive non-specific input) (D) Graph of the maximal displacement during evoked contractions in response to oral and anal stimulation shows significant decreases in the displacement during contractions evoked by anal stimulation in both GCaMP-Ednrb^-/-^ (p=0.0006) and GCaMP-Ednrb^+/-^ mice (p=0.0013) compared to GCaMP-Ednrb^+/+^ mice, but no differences in the displacement of contractions evoked by oral stimulation. A repeated measures two-way ANOVA with Tukey’s test for multiple comparisons was used for statistical analyses in C and D with n=8-13 mice per group.

We then measured and compared evoked motor activity in response to oral and anal electrical stimulation to determine whether changes in neural activation translated into abnormal motility. Interestingly, although there were more myenteric neurons that responded to oral stimulation in GCaMP-Ednrb^-/-^ mice, the resulting displacement during evoked motor activity was not significantly different across genotypes (**Figure 5D**), indicating that the increased neural responses are either from interneurons or dysfunctional motor neurons that do not properly innervate smooth muscle. By contrast, the displacement during motor activity evoked by anal stimulation was significantly lower in magnitude in GCaMP-Ednrb^+/-^ and GCaMP-Ednrb^-/-^ mice compared to GCaMP-Ednrb^+/+^ mice (**Figure 5D**), consistent with the observed decrease in neural responses to anal stimulation.

### HSCR-related changes persisted into adulthood in GCaMP-Ednrb^+/-^ mice

We were surprised that a number of abnormalities identified in GCaMP-Ednrb^-/-^ mice also occurred in GCaMP-Ednrb^+/-^ mice at early post-natal ages, so we next tested whether GCaMP-Ednrb^+/-^ mice exhibited similar defects in adulthood. We measured and compared ENS, ICC and motor activity in proximal colons from adult GCaMP-Ednrb^+/+^ and GCaMP-Ednrb^+/-^ mice at 2-6 months of age.

Consistent with findings from mice at P15, spontaneous activity from myenteric neurons was significantly decreased in adult GCaMP-Ednrb^+/-^ mice (**Figure 6A**). Further, the imbalance in ascending and descending myenteric circuits was also evident in adult GCaMP-Ednrb^+/-^ mice; there were significantly more neurons that specifically responded to oral stimulation and significantly fewer neurons that specifically responded to anal stimulation in GCaMP-Ednrb^+/-^ versus GCaMP-Ednrb^+/+^ mice (**Figure 6B**), indicative of stronger descending and weaker ascending ENS pathways, respectively.

**Figure 6.**
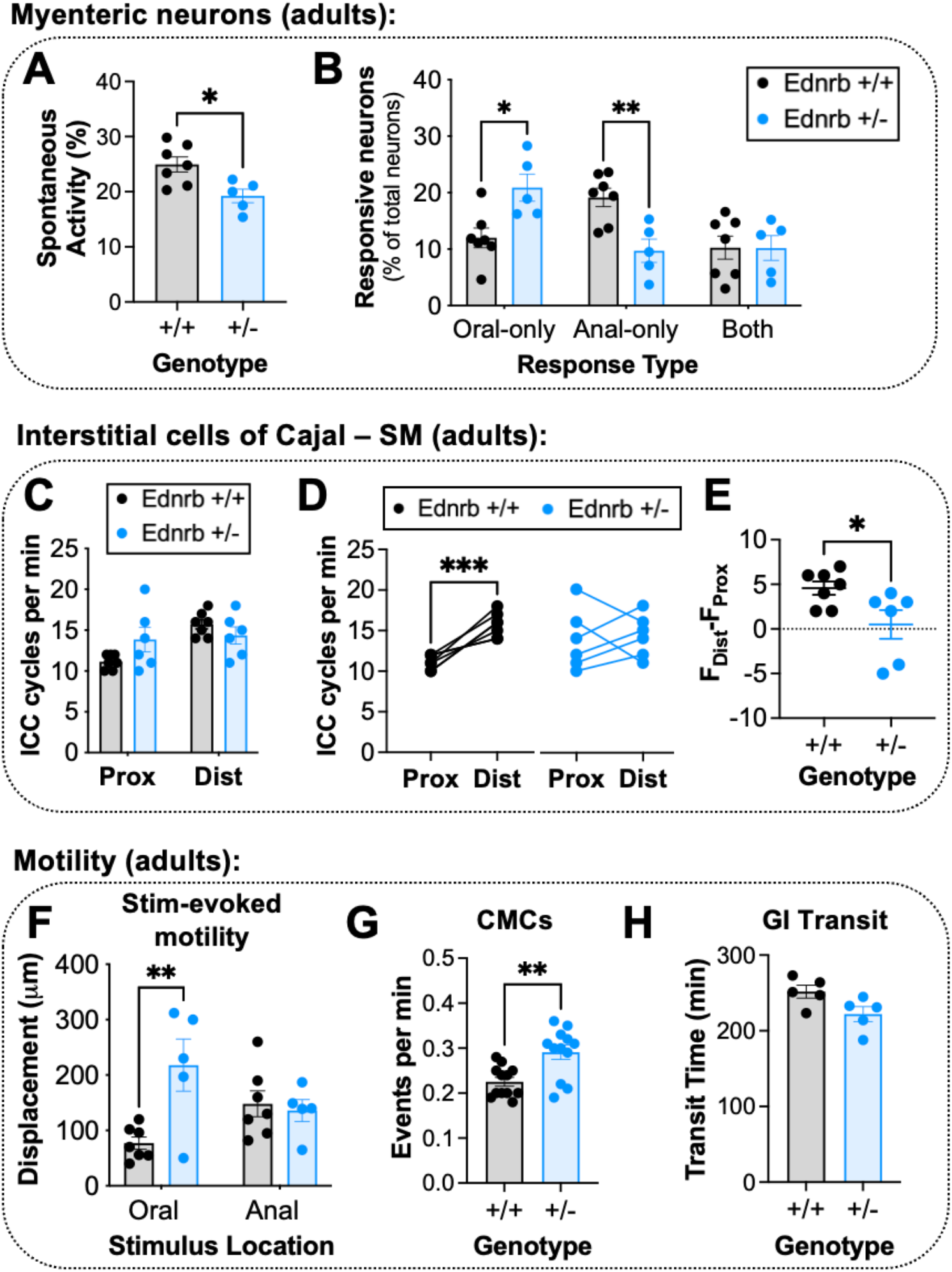
HSCR-related dysfunction in myenteric neuron and ICC activity and motility persists into adulthood in mice with Ednrb het mutations. (A) The average percentage of myenteric neurons in a field of view in the proximal colon that displayed spontaneous activity during 30-sec GCaMP recordings was significantly decreased in adult GCaMP-Ednrb^+/-^ mice compared to GCaMP-Ednrb^+/+^ mice (p=0.0143, n=5-7 mice per group, unpaired Student’s t-test). (B) Adult GCaMP-Ednrb^+/-^ mice had significantly more neurons that specifically responded to oral stimulation (p=0.0119) and significantly fewer neurons that specifically responded to anal stimulation (p=0.0073) compared to age-matched, adult GCaMP-Ednrb^+/+^ mice (n=5-7 mice per group, repeated measures two-way ANOVA with Tukey’s test for multiple comparisons). (C) The frequencies of ICC slow waves in proximal and distal colon were not different between GCaMP-Ednrb^+/-^ and GCaMP-Ednrb^+/+^ mice in adulthood (proximal, p=0.0989 and distal, p=0.5104, n=6-7 mice per group, repeated measures two-way ANOVA with Tukey’s test for multiple comparisons). (D) Similar to P15 mice, the frequency of ICC slow waves were significantly greater in distal colon compared to proximal colon in adult GCaMP-Ednrb^+/+^ mice (p=0.0009, n=7, paired Student’s t-test), but this gradient was absent in adult GCaMP-Ednrb^+/-^ mice (i.e., no differences in frequency between the proximal and distal colon; p=0.7683, n=6, paired Student’s t-test). (E) Plots of ICC slow wave frequency gradient across the colon indicate significant differences between adult GCaMP-Ednrb^+/+^ and GCaMP^+/-^ mice (p=0.0345, n=6-7 mice per group, unpaired Student’s t-test), with the latter exhibiting a lower (or absent) gradient. (F) Tissue displacement during contractions evoked by electrical stimulation oral to the field of view was significantly greater in adult GCaMP-Ednrb^+/-^ mice compared to adult GCaMP-Ednrb^+/+^ mice (p=0.0022, n=5-7 mice per group, repeated measures two-way ANOVA with Tukey’s test for multiple comparisons), but there were no differences in displacement during contractions evoked by anal stimulation. (G) Average frequency of colon motor complexes was significantly higher in adult GCaMP-Ednrb^+/-^ mice compared to GCaMP-Ednrb^+/+^ mice (p=0.0019, n=12, unpaired Student’s t-test). (H) GI total transit time was trending toward a decrease in GCaMP-Ednrb^+/-^ but did not reach significance (p=0.0526, n=5, unpaired Student’s t-test).

Similar to the observations in ICC-SM at P15, adult GCaMP-Ednrb^+/-^ mice also exhibited abnormalities in the frequency gradient of ICC-SM oscillations, but appeared to result from different underlying mechanisms. At P15, the reduced frequency gradient across the colon in GCaMP-Ednrb^+/-^ mice could be attributed to lower frequencies of ICC-SM oscillations in distal colon (**Figure 4**). In adult GCaMP-Ednrb^+/-^ mice, there were no significant differences in the frequency of ICC-SM oscillations in distal colon compared to controls, but there was a strong trend towards an increase in ICC-SM oscillation frequency in proximal colon (**Figure 6C**). The frequency of ICC-SM oscillations in adult GCaMP-Ednrb^+/+^ mice was always higher in distal versus proximal regions from the same colon preparation and created a frequency gradient across the colon (**Figure 6D**). However, this difference between proximal and distal colon was not present in adult GCaMP-Ednrb^+/-^ mice (**Figure 6D**) and the resulting frequency gradient was significantly lower compared to controls (**Figure 6E**). In fact, two GCaMP-Ednrb^+/-^ mice exhibited a reversed frequency gradient indicated by the negative values in Figure 6E. Therefore, whereas decreases in distal colon ICC-SM oscillation frequency contribute to the abnormal frequency gradient in GCaMP-Ednrb^+/-^ mice at P15, increases in proximal colon ICC-SM oscillation frequency appear to contribute to the abnormal frequency gradient in GCaMP-Ednrb^+/-^ mice in adulthood.

Abnormal ENS and ICC activity in proximal colons from adult GCaMP-Ednrb^+/-^ mice correlated to changes in colonic motility behavior. The tissue displacement evoked during oral stimulation was significantly greater in GCaMP-Ednrb^+/-^ mice compared to GCaMP-Ednrb^+/+^ mice, but there were no differences in displacement evoked by anal stimulation (**Figure 6F**). Similar to data regarding ICC-SM, these particular findings in adult mice contrasted with results from P15 mice and indicate that HSCR-associated bowel dysfunction, as well as the contributing mechanisms, may change as ENS circuits mature. The frequency of CMCs was significantly greater in adult GCaMP-Ednrb^+/-^ mice (**Figure 6G**), and while total GI transit time was trending lower in adult GCaMP-Ednrb^+/-^ mice, there were no significant differences between genotypes (**Figure 6H**), suggesting that abnormal motility in adult GCaMP-Ednrb^+/-^ mice was mostly restricted to the colon.

## DISCUSSION

In the present studies, we generated the first HSCR mouse model with GCaMP expression in the ENS and ICC to directly record cellular activity and determine if abnormalities in the ganglionic, proximal colon contribute to post-surgical bowel dysfunction observed in HSCR patients. Based on the significant differences in organization, neurochemical phenotype, spontaneous and evoked activity, and motor patterns in GCaMP-Ednrb^-/-^ mice with HSCR, we conclude that the HSCR proximal bowel, despite the presence of ENS ganglia, is not normal. Dysfunction most likely occurs because the mutant genes that prevent migration of enteric neuron precursors to the distal bowel, thereby causing HSCR-aganglionosis, have additional roles in the developing ENS in proximal bowel regions (e.g., differentiation, synapse formation/pruning). Another possibility is that activity and motility in the distal bowel influence the development of ENS circuits in more proximal regions, and therefore in HSCR, where the distal ENS is absent, the proximal ENS ‘connectome’ does not form correctly. Regardless, our results emphasize that the mere presence of ENS ganglia does not ensure normal bowel function, and additional strategies designed to restore ENS activity in the HSCR proximal bowel should be used in conjuction with surgical removal of distal bowel to reduce long-term dysfunction in HSCR patients.

Although there are a number of HSCR mouse models, we chose to use mice with the loss-of-function mutation in *EDNRB* because homozygosity phenocopies distal bowel aganglionosis used to clinically diagnose HSCR in patients and because it is not lethal until 3-4 weeks of age. Whereas mice with homozygous *EDNRB* mutations have no detectable functional protein, mice with heterozygous *EDNRB* mutations have about 50% protein function, but do not have HSCR-associated aganglionosis, appear phenotypically normal, and live normal lifespans (38). Therefore, we were surprised that many of the abnormalities in post-natal GCaMP-Ednrb^-/-^ mice that model HSCR were also observed in post-natal and adult GCaMP-Ednrb^+/-^ mice. Consistent with our findings, a previous study that used a similar HSCR mouse model with the spotted lethal *EDNRB* mutation (Ednrb^sl^) identified anatomical defects in the distal colon from mice heterozygous for the mutation (Ednrb^s/sl^) that correlated to dysfunction in motor activity, although much less severe than what was observed in mice homozygous for the mutation (Ednrb^sl/sl^) (23). Here, GCaMP-Ednrb^+/-^ mice exhibited hypoganglionosis, alterations in nitrergic myenteric neurons, changes in spontaneous and evoked activity, and signs of dysmotility in the proximal colon. Together, these findings have major implications for studies that have used mice with heterozygous Ednrb mutations as controls. But more importantly, our results suggest HSCR-related mutations that do not produce distal aganglionosis could be the underlying cause of chronic bowel dysfunction in patients for which there is no other clinical explanation (i.e., functional motility disorders, now called disorders of gut brain interactions).

Results from the present study strongly suggest that long-term bowel dysfunction in HSCR patients is due to abnormalities in the proximal colon, but there were some limitations that should be addressed in future experiments. For example, the aganglionic portion of bowel was not removed from post-natal mice with HSCR to mirror patient treatment. Because of the known interactions and long-range projections between proximal and distal colon, it is possible that some of the functional changes we observed were influenced by the presence of an aganglionic distal colon. But as previously mentioned, a number of HSCR-associated changes were also observed in GCaMP-Ednrb^+/-^ mice that have ENS ganglia in distal colon, suggesting that at least some of the dysfunction observed in the proximal colon of Ednrb mutant mice results from decreased Ednrb signaling and would be present whether or not the aganglionic segments were removed. Furthermore, due to the neonatal lethality of GCaMP-Ednrb^-/-^ mice, another limitation of the study is that experiments were performed at an early age before the ENS has completely matured (41, 44–46), so it is unclear whether the observed changes in mice with HSCR persist beyond post-natal ages. It is possible that ENS post-natal developmental processes in HSCR mice (and patients) are simply delayed, and ENS circuits in the proximal colon may become normal, resembling wild-type healthy mice, if given enough time to properly mature. But many of the HSCR-associated changes seen in GCaMP-Ednrb^+/-^ mice at post-natal day 15 were also present in adult GCaMP-Ednrb^+/-^ mice, suggesting that these defects would also persist into adulthood in GCaMP-Ednrb^-/-^ mice. To directly address these limitations and further investigate mechanisms underlying long-term bowel dysfunction in HSCR, future experiments should implement surgical removal of the aganglionic bowel in GCaMP-Ednrb^-/-^ mice allowing survival into adulthood (47).

Based on our results, alternative or additional treatments are needed to improve long-term outcomes for HSCR patients following surgery. There is currently no standard protocol regarding how much bowel to remove relative to the aganglionic segment, and surgeons must guess what is normal versus abnormal bowel based on macroscopic appearance. Typically, surgeons remove the constricted bowel, which corresponds to the aganglionic region, and an additional 1-2 cm. If ENS ganglia are observed in the proximal end of the resected tissue using a neuronal stain, then no additional bowel is removed. Although surgical treatment increases survival, our data indicate that the proximal bowel in HSCR, even at great distances from the aganglionic region, is innervated by an abnormal ENS and likely contributes to post-surgical motility dysfunction in HSCR. However, removing more bowel is not necessarily the best solution and will itself lead to complications and potentially worsen outcomes. Rather, the solution likely involves combining strategies to normalize ENS and/or ICC activity in the abnormal, ganglionic regions that remain after surgery to restore proper function. For example, pacemaker devices could re-entrain ICC slow waves to increase or decrease smooth muscle tension. Extrinsic nerve pathways target the ENS and other non-neuronal cells (e.g., immune, epithelial, and glial cells) to influence various bowel functions (30, 31, 48–54), and therefore nerve stimulation could be utilized to alleviate symptoms associated with dysmotility, inflammation and enterocolitis. Finally, much progress has been made regarding the use of cellular replacement therapies to reestablish functionally-connected enteric neurons using mouse models of enteric neuropathy, including Ednrb mutant mice that model HSCR (55–57). Although not yet a feasible cure for HSCR, cellular therapy could be used in conjunction with surgery to resolve long-term bowel issues caused by the hypoganglionosis in the proximal bowel that remains after surgical treatment. Further, strategies should be developed that increase the differentiation of ENS precursors into excitatory myenteric neurons (e.g., acetylcholine, substance P, and calretinin) to balance the overwhelmingly high proportions of nitrergic (inhibitory) myenteric neurons that populate the proximal bowel in HSCR.

In conclusion, we have shown that the GCaMP-Ednrb mouse model generated in the present study is a valuable translational tool that will continue to advance our understanding of bowel dysfunction in HSCR and should be utilized in future studies to address the gaps in knowledge previously discussed. Here, we identified significant functional defects in the proximal ENS in mice with distal bowel aganglionosis used to clinically diagnose HSCR, and some of these (but not all) were also observed in mice with ‘subclinical’ HSCR-associated mutations (i.e., Ednrb^+/-^), indicating that long-term dysfunction has different underlying causes unrelated to aganglionosis. Comparing and contrasting abnormalities in Ednrb mutant mice (heterozygotes and homozygotes) will help to identify mechanisms of dysfunction and differentiate aspects of ENS development that are dependent on the level of Ednrb expression as well as downstream signaling. Further, because of the ability to assess functional ENS connections, ICC activity, and motor output in real time, the GCaMP-Ednrb model is ideal for optimizing HSCR surgical procedures, measuring the utility of cellular therapies, and testing alternative treatment strategies with the goal of improving the quality of life in patients with HSCR.

## METHODS

### Animals

Mice heterozygous for Ednrb^tm1Ywa^ were purchased from Jax labs (RRID:IMSR_JAX:021933) and bred to a previously established E2a-GCaMP6s mouse line (made by crossing RRID:IMSR_JAX:003724 and RRID:IMSR_JAX:02886) that expresses the GCaMP6s calcium indicator in neuronal and non-neuronal cell types to generate E2a-GCaMP6s mice wild-type (GCaMP-Ednrb^+/+^), heterozygous (GCaMP-Ednrb^+/-^), and homozygous (GCaMP-Ednrb^-/-^) for Ednrb^tm1Ywa^. For post-natal studies, male and female GCaMP-Ednrb^+/+^, GCaMP-Ednrb^+/-^, and GCaMP-Ednrb^-/-^ mice were used at 15 days. For adult studies, male and female GCaMP-Ednrb^+/+^ and GCaMP-Ednrb^+/-^ mice were used at 3-6 months of age. Animals were housed in facilities approved by the Assessment and Accreditation of Laboratory Animal Care (AALAC), with a 12-hour light/dark cycle and free access to water and standard chow. The majority of experiments were completed at the University of Pittsburgh, but a subset of experiments were performed at the Mayo Clinic.

### Immunofluorescence

For immunolabeling in ‘swiss roll’ preparations, full-length colons were dissected, cut along the anti-mesenteric border, pinned flat and fixed for 5 min. The colon was then rolled up starting at the distal end onto the wooden portion of a cotton swab. The rolled-up colon was secured using a dissection pin and fixed for 2 hours. After fixation, the pin was removed and the tissue was cryoprotected in 25% sucrose in 0.01M phosphate-buffered saline (PBS) at 4°C for 24 hours, sectioned (14μm) on a cryostat, and mounted on Superfrost Plus microscope slides (Fisher Scientific, Pittsburgh, PA). Slides were washed three times for 5 min in 0.01M PBS and incubated in 0.01M PBS containing 10% donkey serum, 0.3% Triton, and ANNA-1 (HuC/D; 1:1,000; gift from Dr. Vanda Lennon) antibody for 24 hours at room temperature. Slides were washed in 0.01M PBS, incubated for 90 min at room temperature with a donkey anti-human Cy^TM^ 3 antibody (1:500; Jackson ImmunoResearch, Cat #709-165-149) and coverslipped. Images were collected on the microscope listed above and manually re-stitched to create an image containing the entire length of colon.

For whole-mount colon immunolabeling, full-length colons were dissected, cut along the anti-mesenteric border, pinned flat, and fixed for 2 hours in 4% paraformaldehyde in 0.01M PBS. Tissue was washed in buffer containing 0.1M Tris HCl, 1.5% NaCl, and 0.3% Triton X-100 at pH 7.4 (TNT) three times for 30 min, then incubated overnight at room temperature in TNT containing 20% normal donkey serum and primary antibodies [ANNA-1 (HuC/D; 1:1,000; gift from Dr. Vanda Lennon) and goat anti-nNOS (1:500; Genetex, Cat #GTX89962) or rabbit anti-Calretinin (1:500; Swant, Cat #7697)]. Colons were washed in TNT for 4 hours and incubated overnight at room temperature in TNT containing secondary antibodies at 1:500 [Cy3 anti-human (Cat #709-165-149) and Cy2 anti-goat (Cat #705-225-147) or Cy2 anti-rabbit (Cat #711-225-152) all from Jackson ImmunoResearch]. Sections of colon tissue were placed between two coverslips in a mounting chamber and imaged on a Leica DM4B microscope with Leica EL6000 external light source and Leica DFC7000 T digical camera using 20x objective with appropriate filters. Five fields of view per colon were imaged for analysis.

### GCaMP calcium imaging in the colon

#### Colon preparation

Mice were euthanized with isoflurane and full-length colons removed and placed into a Sylgard-lined dish (Dow Corning, Midland, MI) with circulating carboxygenated (95/5) physiological solution containing in mM: 117.9 NaCl, 4.7 KCl, 25 NaHCO_3_, 1.3 NaH_2_PO_4_, 1.2 MgSO_4_7H_2_O, 2.5 CaCl_2_, 11.1 D-glucose, 2 sodium butyrate, and 20 sodium acetate (all purchased from Sigma-Aldrich, St. Louis, MO). For experiments using electrical stimulation, 1 `μ`M nifedipine (Sigma-Aldrich, St. Louis, MO) was added to reduce spontaneous contractions. Colons were cut open longitudinally, pinned flat using minutien pins with the mucosal side facing down, and transferred to the stage of a Leica DM6 FS upright fluorescence microscope (Leica, Buffalo Grove, IL) equipped with a Photometrics Prime 95B CMOS camera (Roper Scientific, Tucson, AZ) for calcium imaging. Circulating fluid was slowly heated up to 34-37°C, and the colon equilibrated for 20 min prior to calcium imaging.

#### Data collection

GCaMP activity from myenteric neurons and interstitial cells of Cajal (ICC) was recorded in the same preparations; these cell types can easily be distinguished based on morphology, location, and the kinetics of their calcium transients (30). For myenteric neuron activity, only fields of view in the proximal colon (within 20 μm from the cecum, in the area containing large epithelial folds that have a chevron-like appearance) were imaged and compared across GCaMP-Ednrb genotypes. ICC activity was imaged in proximal and distal colon to determine differences across genotypes, colon region, as well as the differences between aganglionic and ganglionic regions in mice with HSCR. GCaMP signals during spontaneous and electrically-evoked activity were imaged with a 20X or 40X objective lens, and images were collected using Metamorph (Molecular Devices, San Jose, CA) at 20 Hz sampling rate with 50 ms exposure and 2 x 2 binning for 30-90 sec in at least 3 fields of view. Electrical stimulation (20 Hz for 1 sec, 100 μs pulse duration) was delivered using a stimulus isolator (A365, World Precision Instruments, Sarasota, FL) and concentric bipolar electrode (FHC Inc, Bowdoin, ME) placed 3 mm oral or anal of the imaging field of view. The order of stimulation location was randomized with 2 min separating each stimulus. All external devices, including the camera and stimulus isolator, were triggered using Spike 2 software via a CED micro1401-4 electronic automatic data processing machine (Cambridge Electronic Design Limited, Milton, Cambridge, UK).

#### Data analysis

Imaging files collected in Metamorph were exported and opened in ImageJ (National Institutes of Health, Bethesda, MD) as multi-TIFF stacks. Movement within the field of view is proportional to changes in tension during contractions (31) and was measured and corrected for using the “Template-Matching, Align Slices in Stack” plugin, which quantifies displacement in the x- and y-axes, representing the circular and longitudinal muscle axes, respectively. Motility traces showing displacement in response to electrical stimulation were created and quantified in Excel (Microsoft, Redmond, WA). After images were corrected for movement, circular regions of interests (ROIs) were placed on cell bodies and mean fluorescence intensity and standard deviation were measured for each ROI. The amplitudes of GCaMP signals were analyzed and quantified by calculating ΔF/F_0_ as % = [(F – F_0_)/F_0_] x 100, where F is the peak fluorescence signal and F_0_ is the mean fluorescence signal at baseline. Only ΔF/F_0_ greater than 4 standard deviations from baseline were considered responses.

### Motility

Motility was assessed in a number of ways: electrically-evoked contractions (described above), spontaneous colonic motor complexes (CMCs), gastrointestinal transit time, and fecal pellet output.

#### Spontaneous CMCs

In the same colon preparations used for GCaMP calcium imaging described above, spontaneous CMCs were measured on the Leica microscope setup but motility was imaged in a field of view for 15-20 minutes at 6.67 Hz sampling rate, 150 ms exposure with a 10X objective lens. A subset of experiments were performed using a Leica M205 FCA equipped with the Leica K8 camera and 1X objective lens. Time-lapse images were collected using the LAS X microscope software at 5 Hz, 200 ms exposure. CMCs were analyzed by tracking the displacement in the field of view and plotting displacement traces, as described in the GCaMP imaging section above. Finally, spontaneous CMCs in empty, closed colon preparations were video-recorded (Sony HDR-CX440), and the frequency of CMCs was determind by manually counting the number of CMCs that propagated at least 1 mm in a period of 20 min. Results were similar across the three experimental conditions.

#### GI Transit Time and Fecal Output

Mice were orally gavaged with 1 ml of 6% carmine red dissolved in methyl cellulose and individually placed back into a clean cage with bedding and access to food and water. After 2 hours, the fecal contents in the cage were collected every 15 min and inspected for the appearance of the red dye. Transit time was calculated as the elapsed time from oral gavage to the appearance of the first red pellet. At the end of the experiment, the total number of fecal pellets was divided by the transit time for a measure of fecal output.

### Statistics

Microscopy images, GCaMP imaging files, and video recordings of colon motility were de-identified and coded for blinded analysis. The genotypes of mice used for GI transit time and fecal output studies were unknown to the experimenter. Data is represented as mean ±SEM, where n=number of mice. Statistical tests were performed in GraphPad Prism (GraphPad Software, San Diego, CA) and included paired and unpaired Student’s t tests, one-way and two-way analysis of variance (ANOVA) with post-hoc tests for multiple comparisons as indicated in the figure legends. Differences were considered significant if P<0.05.

### Study approval

Animal use protocols were approved by the Institutional Animal Care and Use Committees at the University of Pittsburgh and Mayo Clinic.

## AUTHOR CONTRIBUTIONS

BSE: Data acquisition and analyses, manuscript revisions

ESS: Data acquisition and analyses, manuscript revisions

BMD: Conceptualization, experimental design, interpretation of results, manuscript revisions

KMS-E: Conceptualization, experimental design, data acquisition and analyses, interpretation of results, writing and revising manuscript

## ACKNOWLEDGEMENTS

Funding for these studies was provided by the REACH Hirschsprung’s Foundation (KMS-E) and NIH Grants: DK120115 (KMS-E), DK129708 (KMS-E), and DK122798 (BMD). We thank Chris Sullivan for his assistance in breeding and maintaining the mouse colonies used in experiments. We are also grateful to Robert O. Heuckeroth for providing his expertise on HSCR and valued mentorship.

## REFERENCES

1. Heuckeroth RO. Hirschsprung disease - integrating basic science and clinical medicine to improve outcomes. Nat Rev Gastroenterol Hepatol. 2018;15(3):152–67.

2. Bayliss WM, and Starling EH. The movements and the innervation of the large intestine. J Physiol. 1900;26(1-2):107–18.

3. Bayliss WM, and Starling EH. The movements and innervation of the small intestine. J Physiol. 1899;24(2):99–143.

4. Gosain A, and Brinkman AS. Hirschsprung’s associated enterocolitis. Curr Opin Pediatr. 2015;27(3):364–9.

5. Hackam DJ, Reblock KK, Redlinger RE, and Barksdale EM, Jr. Diagnosis and outcome of Hirschsprung’s disease: does age really matter? Pediatr Surg Int. 2004;20(5):319–22.

6. Menezes M, Corbally M, and Puri P. Long-term results of bowel function after treatment for Hirschsprung’s disease: a 29-year review. Pediatr Surg Int. 2006;22(12):987–90.

7. Rescorla FJ, Morrison AM, Engles D, West KW, and Grosfeld JL. Hirschsprung’s disease. Evaluation of mortality and long-term function in 260 cases. Arch Surg. 1992;127(8):934–41; discussion 41-2.

8. Teitelbaum DH, Cilley RE, Sherman NJ, Bliss D, Uitvlugt ND, Renaud EJ, et al. A decade of experience with the primary pull-through for hirschsprung disease in the newborn period: a multicenter analysis of outcomes. Ann Surg. 2000;232(3):372–80.

9. Thakkar HS, Bassett C, Hsu A, Manuele R, Kufeji D, Richards CA, et al. Functional outcomes in Hirschsprung disease: A single institution’s 12-year experience. J Pediatr Surg. 2017;52(2):277–80.

10. Verkuijl SJ, Friedmacher F, Harter PN, Rolle U, and Broens PM. Persistent bowel dysfunction after surgery for Hirschsprung’s disease: A neuropathological perspective. World J Gastrointest Surg. 2021;13(8):822–33.

11. Furness JB. Types of neurons in the enteric nervous system. J Auton Nerv Syst. 2000;81(1-3):87–96.

12. Furness JB, Jones C, Nurgali K, and Clerc N. Intrinsic primary afferent neurons and nerve circuits within the intestine. Prog Neurobiol. 2004;72(2):143–64.

13. Furness JB, Kunze WA, Bertrand PP, Clerc N, and Bornstein JC. Intrinsic primary afferent neurons of the intestine. Prog Neurobiol. 1998;54(1):1–18.

14. Furness JB. The enteric nervous system and neurogastroenterology. Nat Rev Gastroenterol Hepatol. 2012;9(5):286–94.

15. Costa M, Brookes SJ, and Hennig GW. Anatomy and physiology of the enteric nervous system. Gut. 2000;47 Suppl 4:iv15–9; discussion iv26.

16. Sanders KM, Ward SM, and Koh SD. Interstitial cells: regulators of smooth muscle function. Physiol Rev. 2014;94(3):859–907.

17. Wei R, Parsons SP, and Huizinga JD. Network properties of interstitial cells of Cajal affect intestinal pacemaker activity and motor patterns, according to a mathematical model of weakly coupled oscillators. Exp Physiol. 2017;102(3):329–46.

18. Baker SA, Drumm BT, Cobine CA, Keef KD, and Sanders KM. Inhibitory Neural Regulation of the Ca (2+) Transients in Intramuscular Interstitial Cells of Cajal in the Small Intestine. Front Physiol. 2018;9:328.

19. Baker SA, Drumm BT, Skowronek KE, Rembetski BE, Peri LE, Hennig GW, et al. Excitatory Neuronal Responses of Ca(2+) Transients in Interstitial Cells of Cajal in the Small Intestine. eNeuro. 2018;5(2).

20. Baker SA, Leigh WA, Del Valle G, De Yturriaga IF, Ward SM, Cobine CA, et al. Ca(2+) signaling driving pacemaker activity in submucosal interstitial cells of Cajal in the murine colon. Elife. 2021;10.

21. Bhave S, Arciero E, Baker C, Ho WL, Guyer RA, Hotta R, et al. Pan-enteric neuropathy and dysmotility are present in a mouse model of short-segment Hirschsprung disease and may contribute to post-pullthrough morbidity. J Pediatr Surg. 2021;56(2):250–6.

22. Coyle D, O’Donnell AM, Gillick J, and Puri P. Altered neurotransmitter expression profile in the ganglionic bowel in Hirschsprung’s disease. J Pediatr Surg. 2016;51(5):762–9.

23. Ro S, Hwang SJ, Muto M, Jewett WK, and Spencer NJ. Anatomic modifications in the enteric nervous system of piebald mice and physiological consequences to colonic motor activity. Am J Physiol Gastrointest Liver Physiol. 2006;290(4):G710–8.

24. Roberts RR, Bornstein JC, Bergner AJ, and Young HM. Disturbances of colonic motility in mouse models of Hirschsprung’s disease. Am J Physiol Gastrointest Liver Physiol. 2008;294(4):G996–G1008.

25. Zaitoun I, Erickson CS, Barlow AJ, Klein TR, Heneghan AF, Pierre JF, et al. Altered neuronal density and neurotransmitter expression in the ganglionated region of Ednrb null mice: implications for Hirschsprung’s disease. Neurogastroenterol Motil. 2013;25(3):e233–44.

26. Rolle U, Piotrowska AP, Nemeth L, and Puri P. Altered distribution of interstitial cells of Cajal in Hirschsprung disease. Arch Pathol Lab Med. 2002;126(8):928–33.

27. Chen ZH, Zhang YC, Jiang WF, Yang C, Zou GM, Kong Y, et al. Characterization of interstitial Cajal progenitors cells and their changes in Hirschsprung’s disease. PLoS One. 2014;9(1):e86100.

28. Chen X, Meng X, Zhang H, Feng C, Wang B, Li N, et al. Intestinal proinflammatory macrophages induce a phenotypic switch in interstitial cells of Cajal. J Clin Invest. 2020;130(12):6443–56.

29. Nestor-Kalinoski A, Smith-Edwards KM, Meerschaert K, Margiotta JF, Rajwa B, Davis BM, et al. Unique Neural Circuit Connectivity of Mouse Proximal, Middle, and Distal Colon Defines Regional Colonic Motor Patterns. Cell Mol Gastroenterol Hepatol. 2022;13(1):309–37 e3.

30. Smith-Edwards KM, Edwards BS, Wright CM, Schneider S, Meerschaert KA, Ejoh LL, et al. Sympathetic Input to Multiple Cell Types in Mouse and Human Colon Produces Region-Specific Responses. Gastroenterology. 2021;160(4):1208–23 e4.

31. Smith-Edwards KM, Najjar SA, Edwards BS, Howard MJ, Albers KM, and Davis BM. Extrinsic Primary Afferent Neurons Link Visceral Pain to Colon Motility Through a Spinal Reflex in Mice. Gastroenterology. 2019;157(2):522–36 e2.

32. Amiel J, Attie T, Jan D, Pelet A, Edery P, Bidaud C, et al. Heterozygous endothelin receptor B (EDNRB) mutations in isolated Hirschsprung disease. Hum Mol Genet. 1996;5(3):355–7.

33. Auricchio A, Casari G, Staiano A, and Ballabio A. Endothelin-B receptor mutations in patients with isolated Hirschsprung disease from a non-inbred population. Hum Mol Genet. 1996;5(3):351–4.

34. Chakravarti A. Endothelin receptor-mediated signaling in hirschsprung disease. Hum Mol Genet. 1996;5(3):303–7.

35. Kusafuka T, Wang Y, and Puri P. Novel mutations of the endothelin-B receptor gene in isolated patients with Hirschsprung’s disease. Hum Mol Genet. 1996;5(3):347–9.

36. Puffenberger EG, Hosoda K, Washington SS, Nakao K, deWit D, Yanagisawa M, et al. A missense mutation of the endothelin-B receptor gene in multigenic Hirschsprung’s disease. Cell. 1994;79(7):1257–66.

37. Puffenberger EG, Kauffman ER, Bolk S, Matise TC, Washington SS, Angrist M, et al. Identity-by-descent and association mapping of a recessive gene for Hirschsprung disease on human chromosome 13q22. Hum Mol Genet. 1994;3(8):1217–25.

38. Hosoda K, Hammer RE, Richardson JA, Baynash AG, Cheung JC, Giaid A, et al. Targeted and natural (piebald-lethal) mutations of endothelin-B receptor gene produce megacolon associated with spotted coat color in mice. Cell. 1994;79(7):1267–76.

39. Badner JA, Sieber WK, Garver KL, and Chakravarti A. A genetic study of Hirschsprung disease. Am J Hum Genet. 1990;46(3):568–80.

40. Spouge D, and Baird PA. Hirschsprung disease in a large birth cohort. Teratology. 1985;32(2):171–7.

41. Foong JP. Postnatal Development of the Mouse Enteric Nervous System. Adv Exp Med Biol. 2016;891:135–43.

42. Foong JP, Nguyen TV, Furness JB, Bornstein JC, and Young HM. Myenteric neurons of the mouse small intestine undergo significant electrophysiological and morphological changes during postnatal development. J Physiol. 2012;590(10):2375–90.

43. Hao MM, Bornstein JC, Vanden Berghe P, Lomax AE, Young HM, and Foong JP. The emergence of neural activity and its role in the development of the enteric nervous system. Dev Biol. 2013;382(1):365–74.

44. Burns AJ, and Thapar N. Developmental and Postnatal Changes in the Enteric Nervous System. Journal of Pediatric Gastroenterology and Nutrition. 2013;57:S4–S8.

45. de Vries P, Soret R, Suply E, Heloury Y, and Neunlist M. Postnatal development of myenteric neurochemical phenotype and impact on neuromuscular transmission in the rat colon. Am J Physiol Gastrointest Liver Physiol. 2010;299(2):G539–47.

46. Wester T, O’Briain DS, and Puri P. Notable postnatal alterations in the myenteric plexus of normal human bowel. Gut. 1999;44(5):666–74.

47. Zhao L, Cheng Z, Dhall D, Doherty TM, and Frykman PK. A novel corrective pullthrough surgery in a mouse model of Hirschsprung’s disease. J Pediatr Surg. 2009;44(4):759–66.

48. Browning KN, and Travagli RA. Central nervous system control of gastrointestinal motility and secretion and modulation of gastrointestinal functions. Compr Physiol. 2014;4(4):1339–68.

49. De Groat WC, and Krier J. The sacral parasympathetic reflex pathway regulating colonic motility and defaecation in the cat. J Physiol. 1978;276:481–500.

50. Gulbransen BD, Bains JS, and Sharkey KA. Enteric glia are targets of the sympathetic innervation of the myenteric plexus in the guinea pig distal colon. J Neurosci. 2010;30(19):6801–9.

51. Janig W. Integration of gut function by sympathetic reflexes. Baillieres Clin Gastroenterol. 1988;2(1):45–62.

52. Lundgren O. Sympathetic input into the enteric nervous system. Gut. 2000;47 Suppl 4(Suppl 4):iv33–5; discussion iv6.

53. Matheis F, Muller PA, Graves CL, Gabanyi I, Kerner ZJ, Costa-Borges D, et al. Adrenergic Signaling in Muscularis Macrophages Limits Infection-Induced Neuronal Loss. Cell. 2020;180(1):64–78 e16.

54. Spencer N, McCarron SL, and Smith TK. Sympathetic inhibition of ascending and descending interneurones during the peristaltic reflex in the isolated guinea-pig distal colon. J Physiol. 1999;519 Pt 2(Pt 2):539–50.

55. McCann CJ, Cooper JE, Natarajan D, Jevans B, Burnett LE, Burns AJ, et al. Transplantation of enteric nervous system stem cells rescues nitric oxide synthase deficient mouse colon. Nat Commun. 2017;8:15937.

56. Pan W, Rahman AA, Stavely R, Bhave S, Guyer R, Omer M, et al. Schwann Cells in the Aganglionic Colon of Hirschsprung Disease Can Generate Neurons for Regenerative Therapy. Stem Cells Transl Med. 2022.

57. Stamp LA, Gwynne RM, Foong JPP, Lomax AE, Hao MM, Kaplan DI, et al. Optogenetic Demonstration of Functional Innervation of Mouse Colon by Neurons Derived From Transplanted Neural Cells. Gastroenterology. 2017;152(6):1407–18.

